# Altered Regulation of Ia Afferent Input during Voluntary Activity after Human Spinal Cord Injury

**DOI:** 10.1101/2022.05.15.492010

**Authors:** Bing Chen, Monica A. Perez

## Abstract

Sensory input converging on the spinal cord contributes to the control of voluntary activity. Although sensory pathways reorganize following spinal cord injury (SCI), the extent to which sensory input from Ia afferents is regulated during voluntary activity after the injury remains largely unknown. To address this question, we examined the soleus H-reflex and conditioning of the H-reflex by stimulating homonymous and heteronymous nerves [depression of the soleus H-reflex evoked by common peroneal nerve stimulation (D1 inhibition) and the monosynaptic Ia facilitation of the soleus H-reflex evoked by femoral nerve stimulation (FN facilitation)] at rest, and during tonic voluntary activity in humans with and without chronic incomplete SCI. We found that during voluntary activity the soleus H-reflex size increased in both groups compared with rest, but to a lesser extent in SCI participants. Compared with rest, the D1 inhibition decreased during voluntary activity in controls but it was still present in SCI participants. Further, the FN facilitation increased in controls but remained unchanged in SCI participants during voluntary activity compared with rest. Changes in the D1 inhibition and FN facilitation were correlated with changes in the H-reflex during voluntary activity, suggesting an association between outcomes. These findings provide the first demonstration that the regulation of Ia afferent input from homonymous and heteronymous nerves is altered during voluntary in humans with SCI, resulting in lesser facilitatory effect on motor neurons.

## Introduction

Anatomical and physiological studies have shown that sensory input to motor neurons is altered following spinal cord injury (SCI) ^1^. For example, lesions of descending motor tracts in animals result in aberrant sprouting of primary afferents, leading to symptoms of hyperreflexia^2,3^, and prolonged excitatory post-synaptic potentials are observed in motor neurons in response to brief sensory stimulation^4^. In agreement, humans with SCI show prolonged depolarization of motor neurons in response to stimulation of sensory nerves^5^, exaggerated stretch reflexes^6^, and decreased transmission in spinal inhibitory pathways compared with uninjured controls^7-9^. The functional consequences of this altered sensory input conveying to motor neurons following SCI remain largely unknown.

Different mechanisms can contribute to regulate Ia afferent input conveying to motor neurons. For decades, it was thought sensory regulation was accomplished in part through axoaxonic contacts at the terminal of Ia sensory axons from GABAergic neurons that receive innervation from the brain and spinal cord through presynaptic inhibition^10-13^. Recent evidence in animal and humans suggested that facilitation of Ia mediated excitatory postsynaptic potentials (EPSPs) in motor neurons likely occurs when axon nodes are depolarized from the activation of nodal GABAA receptors, which contributes to reduce branch point failure in Ia afferent fibers^14,15^. Thus, GABAergic networks can have both facilitating and inhibitory actions on afferent transmission within the spinal cord at different sites within Ia afferents^16^.

A critical question is how Ia afferent transmission is regulated during voluntary activity after SCI. In uninjured humans, evidence showed that Ia afferent transmission decreases at the onset of a voluntary contraction^17,18^ and during tonic contractions that last for 1-2 min^18,19^ compared to rest and it changes according to the task requirements^20,21^. Following SCI, descending motor pathways converging onto GABAergic interneurons thought to contribute to regulate Ia afferent transmission^22^, are likely altered. Indeed, humans with SCI show lesser corticospinal^23,24^ and H-reflex^25,26^ modulation during voluntary behaviors compared with control participants. We hypothesized that during voluntary activity Ia afferent input exert a lesser facilitatory effect on motor neurons in SCI compared with control participants.

To test our hypothesis, we examined the soleus H-reflex and conditioning of the H-reflex by stimulating homonymous and heteronymous nerves by measuring the depression of the soleus H-reflex evoked by common peroneal nerve (CPN) stimulation (D1 inhibition) and the monosynaptic Ia facilitation of the soleus H-reflex evoked by femoral nerve (FN) stimulation (FN facilitation).

## Materials and methods

### Subjects

Twenty individuals with SCI (50.7±17.3 yr, 3 women) and 20 control subjects (41.5±13.6 yr, 6 women) participated in the study. All subjects were provided written consent to experimental procedures, which were approved by the local ethics committee at Northwestern University (IRB protocol #STU00209996). Participants with SCI had a chronic injury (≥1 year) and were classified using the International Standards for Neurological Classification of Spinal Cord Injury (ISNCSCI) as having a C2-T12 SCI. Five out of the 20 subjects were categorized by the American Spinal Cord Injury Impairment Scale (AIS) as AIS C and the remaining 15 subjects were classified as AIS D. Nine individuals with SCI were currently taking anti-spastic medication (baclofen and/or tizanidine and/or gabapentin; Table 1) at the time of enrollment.

**Table 1.** SCI participants.

| ID | Age<br>(yrs) | Gender | AIS | MAS | Level | Time post<br>injury<br>(yrs) | Medication |
| --- | --- | --- | --- | --- | --- | --- | --- |
| 1 | 61 | F | D | 1 | T8 | 19 | NO |
| 2 | 48 | M | D | 3 | C4 | 12 | BAC |
| 3 | 56 | M | D | 2 | C4 | 9 | GAB |
| 4 | 41 | M | D | 2 | C3 | 2 | BAC,GAB |
| 5 | 46 | M | D | 3 | C5 | 8 | BAC,GAB |
| 6 | 30 | M | C | 1 | C1 | 8 | NO |
| 7 | 37 | M | D | 4 | C7 | 10 | NO |
| 8 | 71 | M | D | 3 | T10 | 6 | NO |
| 9 | 73 | M | C | 1 | C5 | 8 | NO |
| 10 | 38 | M | D | 2 | C5 | 22 | NO |
| 11 | 33 | M | D | 2 | C5 | 8 | NO |
| 12 | 64 | M | C | 2 | C3 | 40 | BAC,GAB |
| 13 | 42 | M | D | 1 | C5 | 11 | NO |
| 14 | 63 | M | C | 1 | C5 | 6 | TIZ |
| 15 | 59 | M | D | 1 | C7 | 11 | BAC |
| 16 | 28 | M | D | 4 | C4 | 1 | BAC |
| 17 | 19 | M | D | 2 | C5 | 2 | NO |
| 18 | 80 | M | D | 1 | C4 | 9 | NO |
| 19 | 58 | F | C | 1 | C3 | 5 | GAB |
| 20 | 72 | M | D | 0 | C3 | 12 | NO |

M = male; F = female; AIS = American Spinal Injury Association impairment scale; BAC = baclofen; GAB = gabapentin; DIA = diazepam; TIZ = tizanidine

These participants were asked to stop anti-spastic medication on the day of testing (at least 12 hours since last dosage). Spasticity was assessed using the Modified Ashworth Scale (MAS). All the SCI and controls participated in the H-reflex experiment, 6 of 20 controls were not able to come back to the lab for the conditioning H-reflex experiments. Sample size was estimated using an effect size (η^2^p=0.3) calculated from the F-statistic (F_1,38_=15.6, p<0.001) for the significant GROUP × CONTRACTION interaction that identified a lesser increase in H-reflex size in SCI compared with control subjects during voluntary contraction. With a power of 0.95 and α of 0.05, 36 participants were considered sufficient in a repeated measures ANOVA (G*Power 3.1.9.7).

### Electromyographic recordings

Electromyography (EMG) was recorded from the soleus and tibialis anterior muscle of the right side in control subjects and from the leg with the higher MAS score in individuals with SCI through bipolar surface electrodes (inter-electrode distance, 2 cm) placed over the belly of the muscle below the gastrocnemius muscles (AG-AgCl, 10-mm diameter). EMG signals were amplified, filtered (20-1,000 Hz), and sampled at 2 kHz for both online detection (3^rd^ order Butterworth, 5-150 Hz band pass filtered and rectified) and offline analysis (CED 1401 with signal software, Cambridge Electronic design, Cambridge, UK).

### Experimental setup

During all testing procedures, subjects were seated comfortably in a custom armchair with both legs placed on a custom platform with the hip (∼120°) and knee (∼160°) flexed and the ankle restrained by straps in ∼110° of plantarflexion. At the start of the experiment, participants performed 3 brief maximal voluntary contractions (MVCs) of 3–5 s into plantarflexion, separated by 30 s. The maximal mean EMG activity in the soleus muscle was measure over a period of 1 s on the rectified response generated during each MVC and the highest value of the three trials was used. The soleus H-reflex (Figure 1, see methods below) was tested at rest and during 30% (controls=29.8±7.1% of MVC and SCI=33.2±4.0% of MVC, p=0.2) of MVC into plantarflexion. Because MVCs were lower in SCI compared with control participants (controls=0.3±0.1 mV and SCI=0.1±0.1mV, p<0.001) we conducted additional experiments in control subjects (n=10) in which we matched the absolute EMG level exerted by SCI participants in all conditions. We also measured Ia afferent transmission (by testing the D1 inhibition and the FN facilitation; Figure 1) at rest and during 30% of MVC. Mean rectified EMG activity in the soleus muscle was shown online on a computer screen located in front of the participants by using Signal software to ensure that individuals were able to match EMG activity during all tasks.

**Figure 1.**
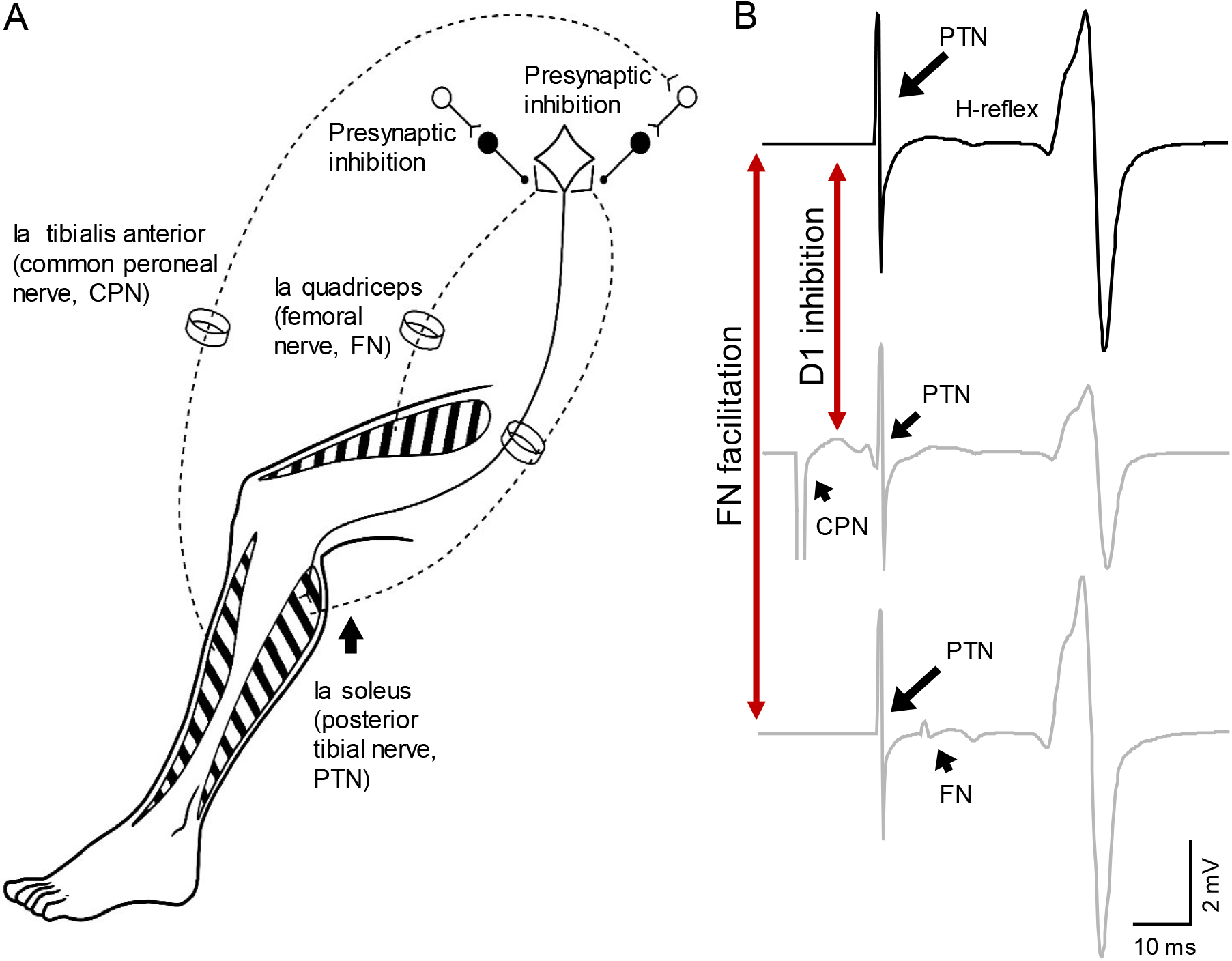
Experimental set-up. (A) Schematic representation of afferent fibers and motor neurons stimulated during our procedures. The soleus reflex was evoked by electrical stimulation of Ia afferents on the posterior tibial nerve (PTN). We assessed presynaptic inhibition by measuring the depression of the soleus H-reflex evoked by stimulating Ia afferents on the common peroneal nerve (CPN) (referred as ‘D1 inhibition’) and the monosynaptic Ia facilitation of the soleus H-reflex evoked by stimulating Ia afferents on femoral nerve (referred as ‘FN facilitation’) at rest, and during tonic voluntary activity. (B) Representative traces showing the soleus H-reflex evoked by PNT stimulation, the D1 inhibition evoked by stimulation of the CPN preceding the PTN at a conditioning-test interval of 15 ms, and the FN facilitation evoked by stimulation of the FN after the PTN at a conditioning-test interval of -8 ms (negative value of the interval indicates that the stimuli to the PTN precedes the FN stimuli).

### Soleus H-reflex

The soleus H-reflex was elicited by using electrical stimulation with the cathode positioned over the posterior tibial nerve in the popliteal fossa and the anode positioned above the patella using a constant-current stimulator (1 ms rectangular electrical stimulus, 0.25 Hz; model DS7A, Digitimer, Hertfordshire, UK). The reflex response was measured as peak-to-peak amplitude of the non-rectified reflex response recorded from the soleus muscle. The stimulus intensity was increased in steps of 0.05 mA starting below H-reflex threshold and increasing up to supramaximal intensity to measure the maximal motor response (M-max). To ensure that M-max values were reached, the stimulus intensity was increased until a plateau was observed in the M-max (controls=13.2±3.3 mV and SCI=10.3±4.8 mV, p=0.03). The size of the H-reflex was kept at 50% of the maximal H-reflex (H-max; controls=6.2±2.7 mV and SCI=6.5±4.0 mV, p=0.8) at rest and the same intensity was used during voluntary activity. The magnitude of H-reflex during 30% of MVC was expressed as a % of the H-reflex at rest. Twenty reflexes were tested at test and 20 reflexes were tested during 30% of MVC in a randomized order.

### D1 inhibition

The soleus H-reflex (control H-reflex) was conditioned by stimulation of the common peroneal nerve (CPN, conditioned H-reflex). The CPN stimulation elicits a depression of the soleus H-reflex at conditioning-test interval of 8-20 ms (referred to as D1 inhibition^27^). Consistent with previous results^27,28^, we used a conditioning-test interval of 15 ms to assess the D1 inhibition at rest and during 30% of MVC in both controls (n=14) and SCI participants (n=20). The CPN was stimulated (1 ms rectangular electrical stimulus) through bipolar a bar electrode placed over the nerve distal to the neck of the fibula. The goal was to evoke a motor response in the tibialis anterior muscle without a motor response in the peroneal muscles. The intensity of the CPN stimulation was kept at 1.4× motor response threshold (MT) in the tibialis anterior muscle^27^. The MT was defined as the minimal intensity needed to elicit 5 of 10 motor responses in the tibialis anterior muscle of 50 µV above the background. The size of the control H-reflex was kept at 50% of the maximal H-reflex at rest and the same intensity was used during voluntary activity. We found that the magnitude of the D1 inhibition was decreased in SCI (81.2±7.8%) compared with control (69.6±15.4%, p=0.01) subjects at rest. Thus, in an additional control experiment, we tested the D1 inhibition in SCI participants (n=10) using a higher CPN stimulation at 1.5-2×MT while the control H-reflex was kept at 50% of the H-max to elicit a magnitude of D1 inhibition similar to controls (referred to as D1 inhibition adjusted, D1 inhibition_adj_). Fifteen control H-reflexes and 15 conditioned H-reflexes were tested at rest and during 30% of MVC in a randomized order.

### FN facilitation

The soleus H-reflex was tested with (conditioned H-reflex) and without (control H-reflex) stimulation of the FN. The FN elicits a facilitation of the soleus H-reflex (FN facilitation), which is thought to reflect the size of the monosynaptic excitatory postsynaptic potential in the soleus motor neurons evoked by activation of Ia afferents from the quadriceps muscle, and changes in its size are considered to indicate changes in Ia afferent transmission^17^. Thus, both measurements, D1 inhibition and FN facilitation, provide independent information about Ia afferent transmission and help to rule out changes in the recruitment gain of soleus motor neurons^18^. The FN was stimulated through bipolar electrodes with the cathode positioned over the femoral triangle and the anode electrode positioned just below the gluteus maximus muscle. The intensity for stimulating the FN was 5×MT in the quadriceps muscle^29,30^. The onset of facilitation was taken to be the earliest conditioning-test interval at which the conditioned reflex was at least 5% larger than the control reflex to ensure that the conditioning-test interval reflects the size of the monosynaptic excitatory postsynaptic potential in the soleus motor neurons without contamination^31^. Measurements were taken at 0.5-1 ms longer than this interval. The size of the control H-reflex was kept at 50% of the maximal H-reflex at rest and the same intensity was used during 30% of MVC. The facilitation induced by stimulating the FN has an onset at a conditioning-test interval between -7 and -8.5 ms^18,29^ (negative value of the interval means that the control stimulus precedes the conditioning stimulus). In a control experiment, we tested conditioning-test intervals between –6.5 and –9.0 ms (negative value of the interval means that the control stimulus precedes the conditioning stimulus) in controls (n=5) and SCI (n=5) and determined that in both groups the earliest onset of the FN was found at -7.5 ms (p=0.01). Thus, consistent with ours and previous results^18,29^, we used a conditioning-test interval of -8 ms to evaluate the FN facilitation at rest and during 30% of MVC in both controls (n=14) and SCI participants (n=20). The magnitude of the FN facilitation at rest is decreased in SCI compared with control subjects^29^, therefore, in a control experiment we tested the FN facilitation in SCI participants (n=10) using lower FN stimulation at 2-4×MT (referred to as FN facilitation adjusted, FN facilitation_adj_) while the control H-reflex was kept at 50% of the H-max. Fifteen control H-reflexes and 15 conditioned H-reflexes were tested at rest and during 30% of MVC in a randomized order.

### Data analysis

Normal distribution was tested by the Shapiro-Wilk’s test and homogeneity of variances by the Levene’s test. Sphericity was tested using Mauchly’s test. When sphericity was not met, the Greenhouse-Geisser correction was used. Repeated measures ANOVA was used to examine the effect of GROUP (controls and SCI) and CONDITION (rest and 30% of MVC) on background EMG and H-reflex. Repeated measures ANOVA was also used to examine the effect of GROUP and CONDITION on the D1 inhibition, and FN facilitation. Similar analysis was also used to examine the effect of CONDITION on the D1 inhibition_adj_ and the FN facilitation_adj_. A one-way ANOVA was used to examine the effect of GROUP (controls and SCI) on the ratio of H-max and M-max (H/M ratio). Holm-Sidak *post hoc* analysis was used to test for mean pair wise comparisons. Spearman’s correlation coefficient was used to assess association between the size of H-reflex, D1 inhibition, FN facilitation during 30% of MVC. Statistical analysis was conducted using SigmaPlot (Systat Software, Inc, San Jose, CA, USA) and the significance was set at p<0.05. Group data is presented as means±SDs. Effect sizes were reported as η^2^p.

## Results

### EMG

Figure 2A shows raw rectified EMG data in the soleus muscle in representative control and SCI participants. Note that EMG activity increased during voluntary contraction in both participants compared with rest but to a lesser extent in SCI subjects. Repeated measures ANOVA showed an effect of GROUP (F_1,38_=13.7, p=0.001, η^2^p=0.3), CONTRACTION (F_1,38_=143.8, p<0.001, η^2^p=0.8) and in their interaction (F_1,38_=13.7, p=0.001, η^2^p=0.3) on mean rectified EMG activity normalized to the MVC. *Post-hoc* analysis showed that EMG activity increased at 30% of MVC (controls, p<0.001 and SCI, p<0.001) compared with rest in both groups. One-way ANOVA showed an effect of GROUP on the M-max (controls=13.2±3.3 mV and SCI=10.3±4.8 mV, p=0.03, η^2^p=0.2) but not the H-max (controls=6.2±2.7 mV and SCI=6.5±4.0 mV, p=0.8) in the soleus muscle. We found a Group effect on H/M ratio (controls=48.5±19.7%, SCI=62.0±19.1%, p=0.03, η^2^p=0.2). Note that there was no difference in the activation of the tibialis anterior muscle between controls and SCI participants during 30% of plantarflexion MVC (controls=7.3±3.6% of tibialis anterior MVC and SCI=9.2±10.6% of tibialis anterior MVC, p=0.2).

**Figure 2.**
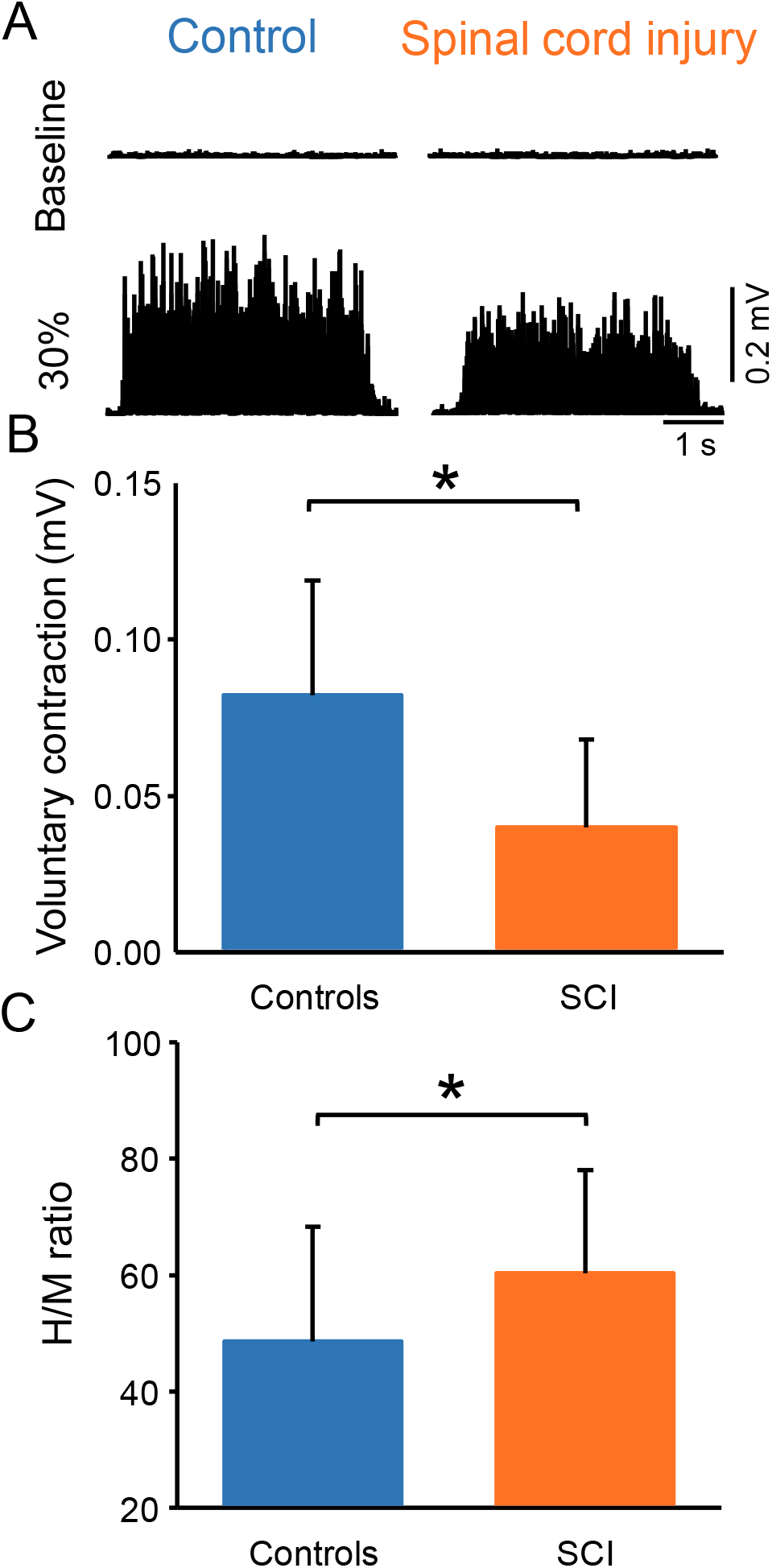
Voluntary contraction and maximal H-reflex and maximal motor response (M-max) ratio (H/M ratio). (A) Electromyographic (EMG) traces tested at rest and during 30% of maximal voluntary contraction (MVC) with the soleus muscle in a control and in a SCI participant. (B) The bar graph shows group data of 30% of MVC. The abscissa shows the groups tested (controls=blue bar, SCI=orange bar) and the ordinate shows the MVC (in millivolts). (C) Bar graph shows the group data on the H/M ratio. The abscissa shows the groups tested (controls=blue bar, SCI=orange bar) and the ordinate shows the H/M ratio. *p<0.05.

### Soleus H-reflex

Figures 3A illustrate raw traces showing the soleus H-reflex in a control and a SCI participant. Note that the H-reflex increased in both participants during voluntary activity comparted with rest but to a lesser extent in the individual with SCI. Repeated measures ANOVA showed an effect of GROUP (F_1,38_=15.6, p<0.001, η^2^p=0.3), CONTRACTION (F_1,28_=100.3, p<0.001, η^2^p=0.7) and in their interaction (F_1,38_=15.6, p<0.001, η^2^p=0.3) on the H-reflex size. *Post-hoc* analysis showed that the H-reflex was larger during 30% of MVC (controls=245.6±88.7%, p<0.001; SCI=163.2±23.2%, p<0.001) compared to rest in both groups. Additionally, the H-reflex size increased to a lesser extent in SCI compared to control subjects at 30% of MVC (p<0.001).

**Figure 3.**
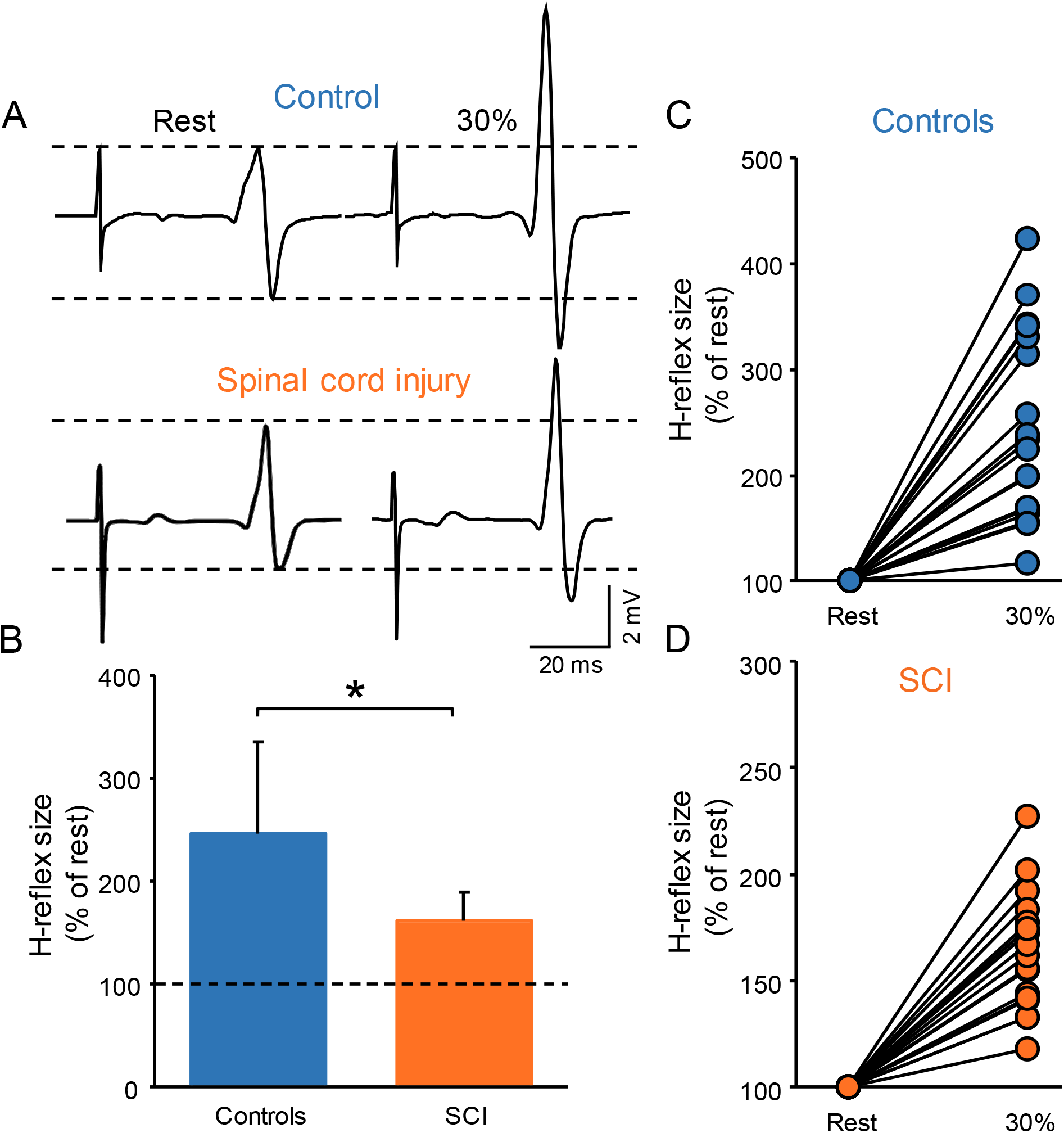
Soleus H-reflex. (A) Representative EMG traces showing the soleus H-reflex tested at rest and during 30% of MVC in a control and in a SCI participant. (B) Graph shows the group H-reflex data. The abscissa shows the groups tested (controls=blue bar, SCI=orange bar) and the ordinate shows the H-reflex size during 30% of MVC expressed as a % of the H-reflex size at rest. Graphs show individual H-reflex data in controls (C) and SCI (D) participants. The abscissa shows the conditions tested (rest, 30% of MVC) and the ordinate shows the H-reflex size during 30% of MVC expressed as a % of the H-reflex size at rest. *p<0.05.

We conducted an additional control experiment where we matched absolute EMG level across groups during 30% of MVC by asking control participants to perform the similar EMG activity as SCI participants. Repeated measures ANOVA showed an effect of GROUP (F_1,18_=6.0, p=0.02, η^2^p=0.3), CONTRACTION (F_1,18_=52.4, p<0.001, η^2^p=0.7) and in their interaction (F_1,18_=6.1, p=0.02, η^2^p=0.3) on H-reflex amplitude. *Post-hoc* analysis showed that the H-reflex was larger during 30% of MVC (controls=218.9±73.7%, p<0.001; SCI=158.12±23.1%, p<0.001) compared to rest in both groups. Note that the increases in H-reflex size were lesser in SCI compared with control at 30% of MVC (p=0.02).

### D1 inhibition

Figures 4A and B illustrate raw traces showing the D1 inhibition measured in a representative control and SCI participant. Compared with rest, it shows that the D1 inhibition was abolished during 30% of MVC in the control participant while the D1 inhibition remained unchanged in the SCI participant. Repeated measures ANOVA showed an effect of GROUP (F_1,32_=6.4, p=0.01, η^2^p=0.1), CONTRACTION (F_1,32_=36.6, p<0.001, η^2^p=0.6) and in their interaction (F_1,32_=20.6, p<0.001, η^2^p=0.7) on the D1 inhibition. *Post-hoc* analysis showed that the D1 inhibition decreased during 30% of MVC in controls (rest=69.6±15.4%, 30% of MVC=100.5±7.2%, p<0.001) but not in SCI (rest=81.2±7.8%, 30% of MVC=80.2±6.6%, p=0.5; Figure 4C) participants. Because D1 inhibition at rest is decreased in SCI (81.2±7.8%) compared with control (69.6±15.4%, p=0.02) participants, we tested the D1 inhibition in a subgroup of SCI participants (n=10) by adjusting to magnitude of the D1 inhibition to match the values obtained in control subjects (D1 inhibition_adj_, see Methods). Here, we found that the D1 inhibition_adj_ decreased during 30% of MVC compared with rest in SCI participants (rest=72.9±6.1%, 30% of MVC=81.1±8.2%, p<0.001; Figure 4C-D).

**Figure 4.**
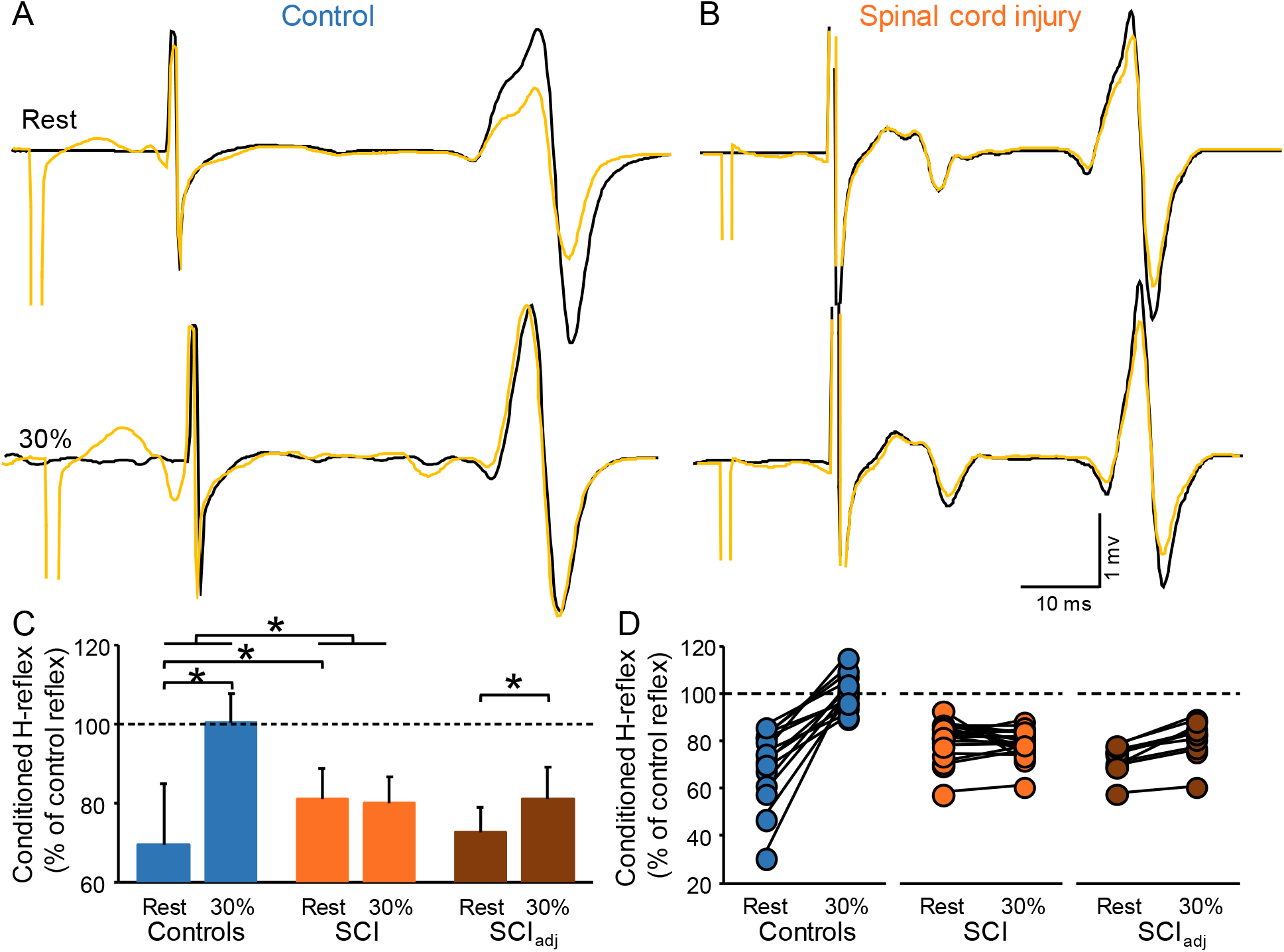
D1 inhibition. Representative traces showing the H-reflex (control reflex, in black) and the H-reflex conditioned by CPN stimulation (conditioned H-reflex, in yellow) tested at rest and during 30% of MVC in a control (A) and in a SCI (B) participant. The bar graph shows the conditioned H-reflex normalized to the control H-reflex in both groups (C). The abscissa shows the groups tested at rest and during 30% of MVC (controls=blue bars, SCI=orange bars, SCI_adj_=brown bars). Note that here the SCI_adj_ condition refers to testing of the D1 inhibition_adj_. The ordinate shows the size of conditioned H-reflex expressed as a % of the control H-reflex (use to assess the D1 inhibition). Data from individual subjects (D) showing the conditioned H-reflex normalized to the control H-reflex in all groups tested (controls=blue circles, SCI=orange circles, SCI_adj_=brown circles). *p<0.05.

### FN facilitation

Figures 5A and B illustrate raw traces showing the FN facilitation measured in a control and in a SCI participant. We found that the FN facilitation increased in the control but not in the SCI participant during 30% of MVC compared with rest. Repeated measures ANOVA showed an effect of GROUP (F_1,32_=4.8, p=0.04, η^2^p=0.1), CONTRACTION (F_1,32_=30.8, p<0.001, η ^2^p=0.5) and in their interaction (F_1,30_=51.5, p<0.001, η^2^p=0.6). *Post-hoc* analysis showed that the FN facilitation increased during 30% of MVC in controls (rest=111.0±7.1%, 30% of MVC=126.3±8.2%, p<0.001) but not in SCI (rest=119.2±9.3%, 30% of MVC=117.2±7.6%, p=0.16; Figure 5C) participants. The FN facilitation at rest was increased in SCI (119.2±9.3%) compared with control (111.0±7.1%, p=0.007) participants. Thus, we tested the FN facilitation in a subgroup of SCI participants (n=10) by adjusting to magnitude of the intensity of the conditioning pulse to match the level of FN facilitation obtained in control participants (FN facilitation_adj_, see Methods). We found that the FN facilitation_adj_ increased during 30% of MVC and rest in SCI participants (rest=113.2±6.2%, 30% of MVC=120.2±11.1%, p=0.01; Figure 5C-D).

**Figure 5.**
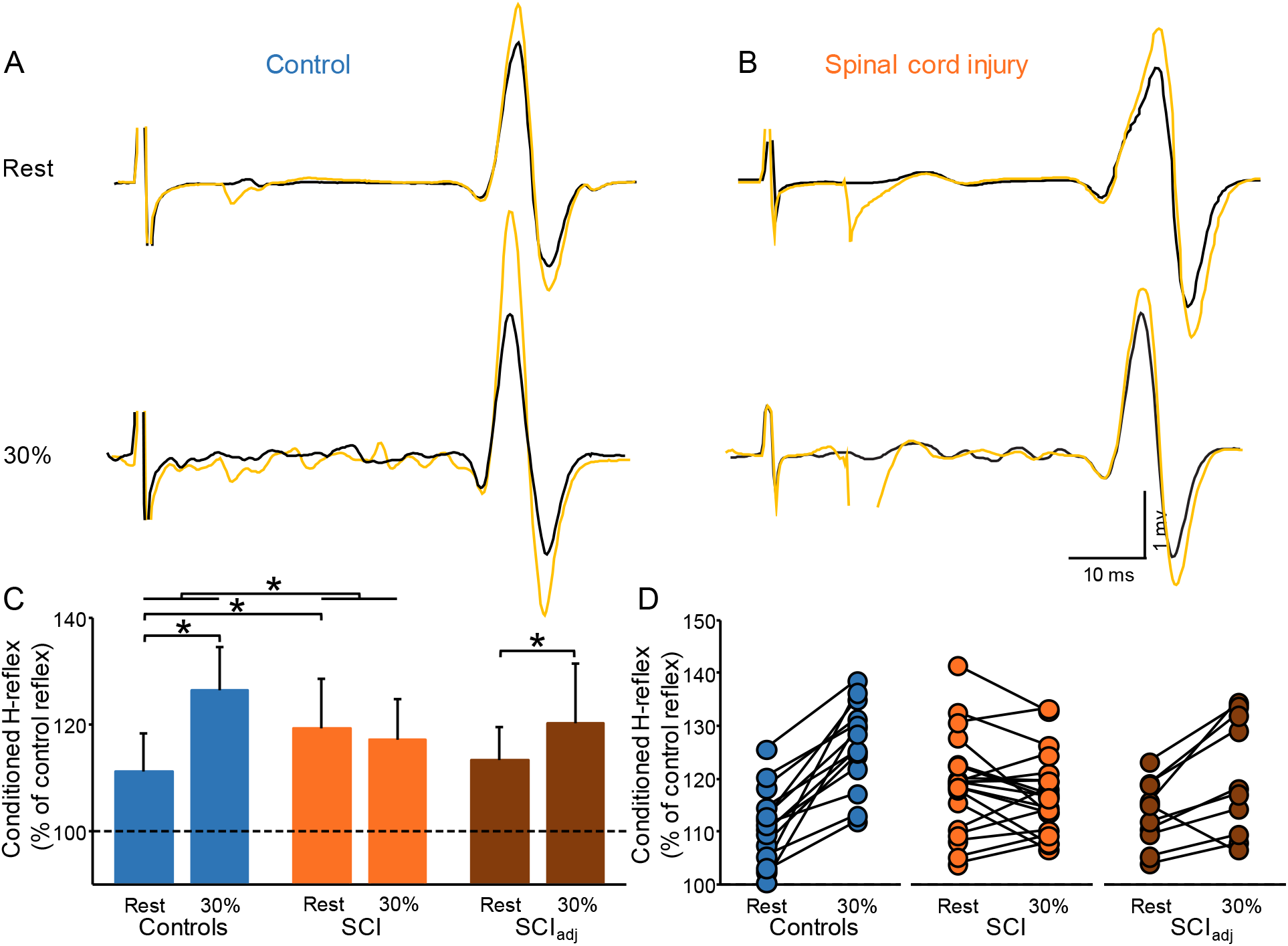
FN facilitation. Representative traces showing the H-reflex (control reflex, in black) and the H-reflex conditioned by FN stimulation (conditioned H-reflex, in yellow) tested at rest and during 30% of MVC in a control (A) and in a SCI (B) participant. The bar graph shows the conditioned H-reflex normalized to control H-reflex in both groups (C). The abscissa shows the groups tested at rest and during 30% of MVC (controls=blue bars, SCI=orange bars, SCI_adj_=brown bars). Note that here the SCI_adj_ condition refers to the testing of the FN facilitation_adj_. The ordinate shows the size of the conditioned H-reflex expressed as a % of the control H-reflex (use to assess the FN facilitation). Data from individual subjects (D) showing the conditioned H-reflex normalized to the control H-reflex in all groups tested (controls=blue circles, SCI=orange circles, SCI_adj_=brown circles). *p<0.05.

### Correlation

Figure 6 shows the correlation between the size of H-reflex, D1 inhibition, and FN facilitation during 30% of MVC compared with rest. We found that the H-reflex size positively correlated with the D1 inhibition (r=0.6, p<0.001) and the FN facilitation (r=0.4, p=0.02) during 30% of MVC compared with rest. The D1 inhibition positively correlated with and the FN facilitation (r=0.7, p<0.001) during 30% of MVC compared with rest.

**Figure 6.**
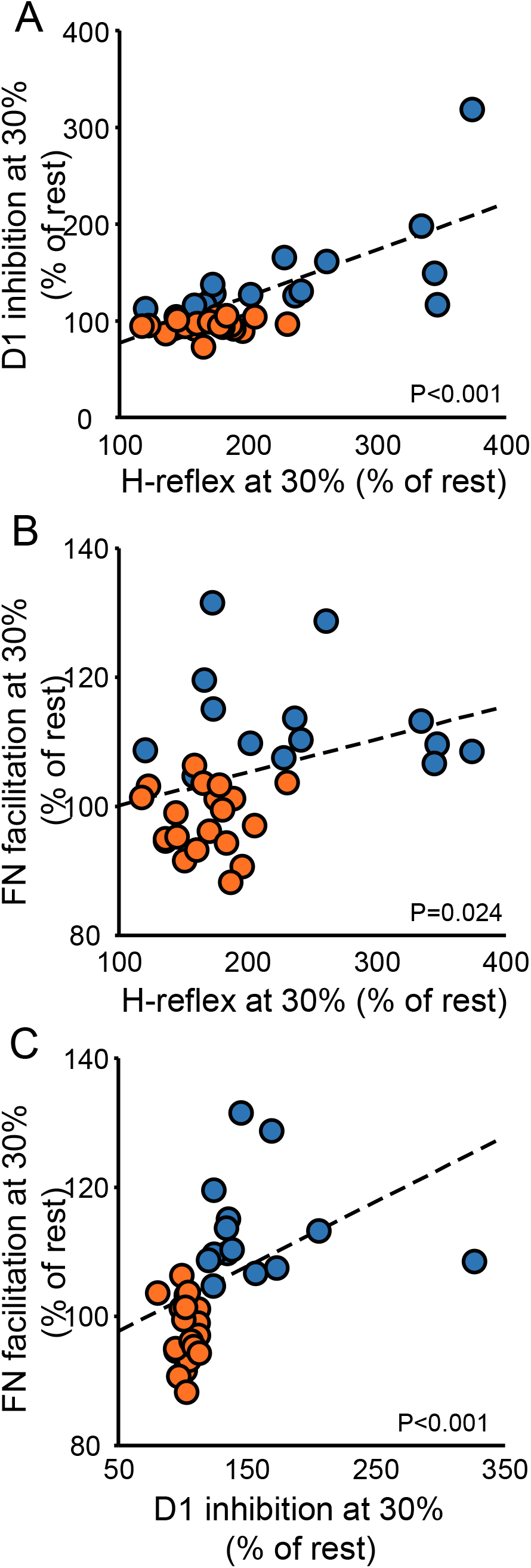
Correlation. Graphs show individual data from controls (blue circles) and SCI (orange circles) participants. The abscissa shows the size of the H-reflex during 30% of MVC expressed as a % of the H-reflex tested at rest (A and B) and the ordinate shows the D1 inhibition during 30% of MVC expressed as a % of the D1 inhibition tested at rest (A), the FN facilitation during 30% of MVC expressed as a % of the FN facilitation tested at rest (B). Data from individual subjects (C) showing the correlation between the FN facilitation during 30% of MVC expressed as a % of the FN facilitation tested at rest (C, abscissa) and the D1 inhibition during 30% of MVC expressed as a % of the D1 inhibition tested at rest (C, ordinate). *p<0.05.

## Discussion

Our findings support the hypothesis that during voluntary activity the regulation of Ia afferent input to motor neurons is altered following SCI. We found that the size of the H-reflex in the soleus muscle increases in controls and SCI participants during voluntary activity but to a lesser extent in people with SCI. Two observations suggest that altered regulation of inputs from homonymous and heteronymous nerves during voluntary might contribute to these results. First, we found that the D1 inhibition decreased in controls but it was still present in SCI participants during voluntary contraction compared with rest. Second, we found that the FN facilitation increased in controls but not in SCI participants during contraction compared with rest. Changes in the D1 inhibition and the FN facilitation correlated with changes in the H-reflex size during voluntary contraction. Together, our findings suggest that during voluntary activity Ia afferent input from homonymous and heteronymous nerves exert a lesser facilitatory effect on motor neurons in humans with SCI compared with control subjects.

### Regulation of Ia afferent input after SCI

Although there have been multiple studies in animals and humans showing that sensory input conveying to motor neurons is altered following SCI^1^ the functional consequences of these changes remain largely unknown. Here, for the first time, we examined the regulation of Ia afferent input during voluntary activity in humans with chronic incomplete SCI. Studies have used H-reflex conditioning paradigms in humans to make inferences about the regulation of Ia afferent input^17-19,32^. For example, the depression of the soleus H-reflex evoked by CPN stimulation (referred here as D1 inhibition) is thought to be caused by presynaptic inhibition at the terminal of Ia afferents on soleus motor neurons^27^ and the FN facilitation is thought to reflect the size of the monosynaptic excitatory postsynaptic potential in the soleus motor neurons evoked by activation of Ia afferents from the quadriceps muscle^32^. Both outcomes likely provide independent information about Ia afferent transmission that help to rule out changes in the recruitment gain of soleus motor neurons^18^. We found that the D1 inhibition was decreased during voluntary activity in controls subjects but still present in SCI participants compared with rest. We also found that the FN facilitation was increased in control subjects during voluntary activity but not in SCI participants when compared with rest. This is consistent with studies showing that, during a ramp up and hold tonic contraction, the FN facilitation decreased at the onset of a voluntary contraction^19,32^ and during a tonic contraction lasting for 1-2 min^18^ as performed in our study. This is also consistent with findings showing that vibratory inhibition of the soleus H reflex decreases during tonic voluntary contraction^33^. These results were previously interpreted as a decreased in presynaptic inhibition during a voluntary contraction. For decades, it was thought that sensory regulation was accomplished in part through axoaxonic contacts at the terminal of Ia sensory axons through presynaptic inhibition^10-12,34^. However, recent evidence in animal and humans suggest that facilitation of Ia mediated EPSPs in motor neurons likely occurs when axon nodes are depolarized from the activation of nodal GABA_A_ receptors^14,15^. Thus, it is possible that the regulation of Ia afferent input tested in our study occurs at different sites within the Ia afferent fiber. We can also not exclude the possibility that the suppression of the H-reflex evoked by conditioning of a homonyomous nerve is related, at least in part, to post-activation depression or any direct effect on the soleus motoneurons^16^. Regardless of the site of Ia afferent regulation, together, our findings suggest that proprioceptive input from homonymous and heteronymous nerves exert a lesser facilitatory effect on motor neurons after SCI.

Both peripheral^35,36^ and central^37-39^ mechanisms have been shown to contribute to regulate Ia sensory transmission. GABAergic neurons contributing to regulate Ia sensory transmission receive innervation from the brain and spinal cord^13^. At rest, decreases in the D1 inhibition and increases in the FN facilitation in people with SCI compared with control participants could be related to decreased input from descending motor pathways. This is supported by multiple demonstrations that activation of descending motor pathways, including the corticospinal pathway, after the injury have a higher threshold resulting in the use of higher stimulus intensities in SCI compared with control participants^23,40,41^. The same explanation can apply during voluntary activity. Our participants had incomplete injuries and were able to perform voluntary activity. Evidence showed that corticospinal excitability increases in controls and SCI participants during voluntary activity but just to a lesser extent in SCI participants^23,24,42^. Thus, a possibility is that the lesser facilitatory effect of Ia afferent input on motor neurons after SCI during voluntary activity reflects altered contribution from descending motor pathways. Note that when the D1 inhibition and FN facilitation were tested at matching levels between groups, we observed a small but significant decreased in the D1 inhibition and increase in the FN facilitation during voluntary activity, suggesting that descending motor pathways may also contribute to these results. This is consistent with our findings showing that H-reflex size increased during voluntary activity in controls and SCI participants but to a lesser extent in people with SCI. Similarly, evidence showed that during voluntary activity, motor neuron excitability (as measured by F waves) increases in people with SCI but to a lesser extent than in control participants ^43^. Stretch reflexes^44^ and H-reflexes^29,45^ have been reported to increase to a lesser extent or not at all during voluntary activity in people with SCI compared with control subjects. Then, how do we explain the facilitatory effect of Ia afferent input on motor neurons in control participants during voluntary activity? A possibility is that this is, in part, related to descending inhibition of GABAergic interneurons, which overrides the suppression originated from peripheral sources during a voluntary contraction. Hence, in SCI participants, a lesser facilitatory effect of Ia afferent input on motor neurons might be present during voluntary activity because of the abnormal and/or decreased contribution from descending pathways, which might not be strong enough to override peripheral sources.

### Functional consequences

During a voluntary contraction, the “excitability” of motor neurons increases. Thus, regulation of Ia afferent input to motor neurons can have implications for the generation of motor output. Indeed, evidence showed that presynaptic regulation of spinal sensory feedback contributes to ensure smooth execution of movement^46^ and motor stability^47^. In intact humans, lesser facilitatory effect of Ia afferent input on motor neurons have been related to the optimization needed during the learning of motor skills^28^. We propose that the lesser facilitatory effect of Ia afferent input on motor neurons during voluntary activity in SCI compared with controls reflects contribution to regulate the ongoing voluntary activity. Since after SCI prolonged excitatory post-synaptic potentials are observed in motor neurons in response to even brief sensory stimulation^4^, a lesser facilitatory effect of Ia afferent input on motor neurons at the spinal level in SCI participants might help to control the ongoing voluntary activity. In controls, the greater facilitatory effect of Ia afferent input onto motor neurons might be functionally appropriate during the tonic voluntary activity that we tested, but this might also change to a different extent during performance of more skilled motor behaviors.

